## Supplemental Figures for "The epigenomic landscape of single vascular cells reflects developmental origin and identifies disease risk loci"

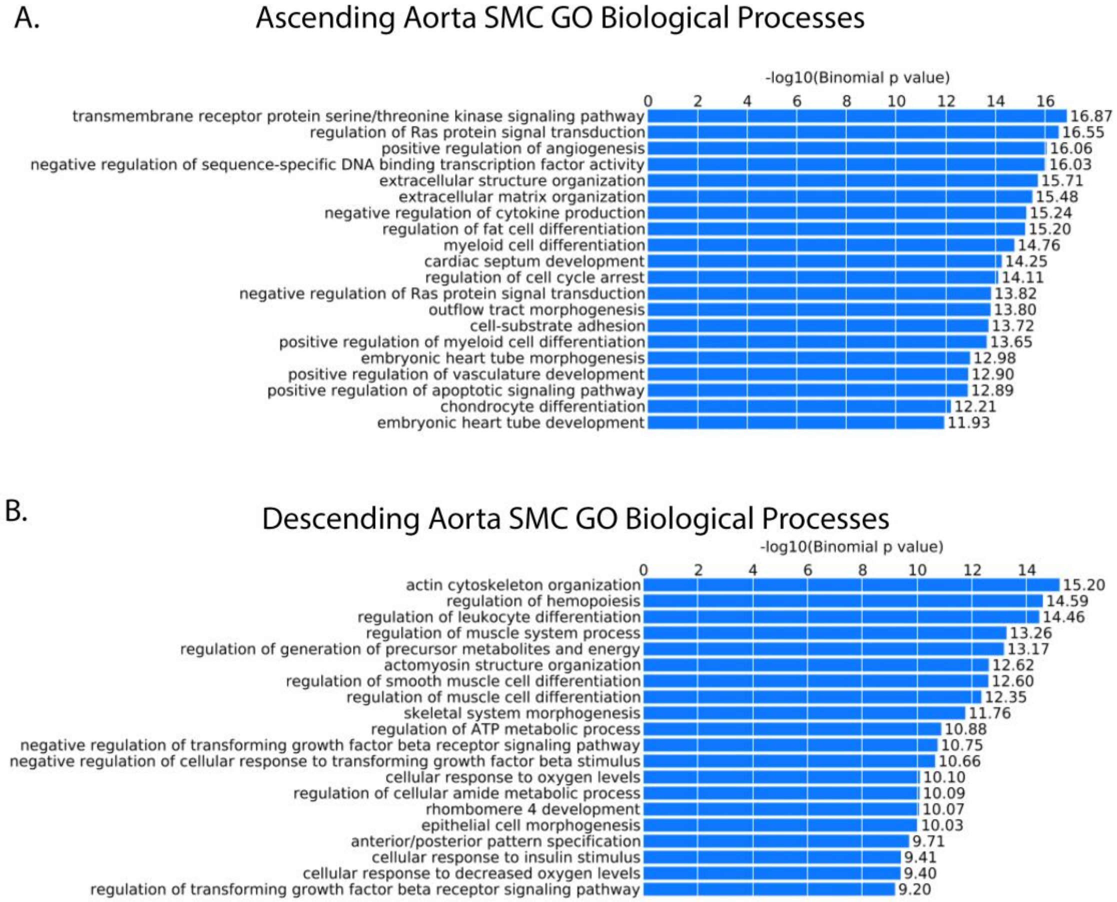


**Supplemental Figure 1.** Top biological processes from GREAT for peaks marking SMCs isolated from the ascending aorta (A) and descending aorta (B).

A.

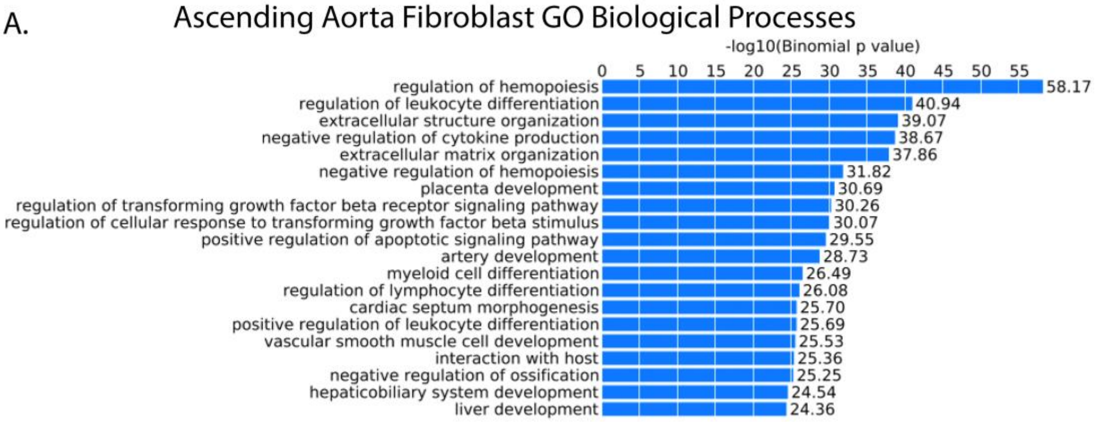


B.

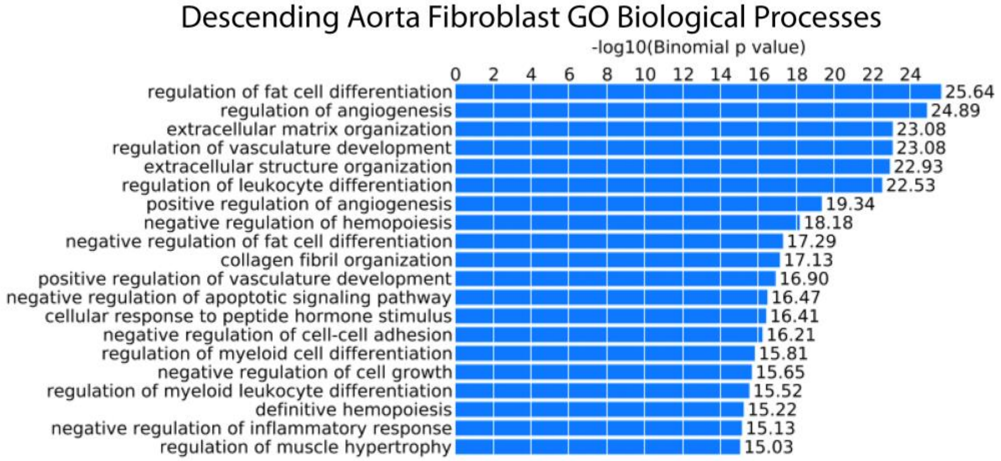


**Supplemental Figure 2.** Top biological processes from GREAT for peaks marking fibroblasts isolated from the ascending aorta (A) and descending aorta (B).

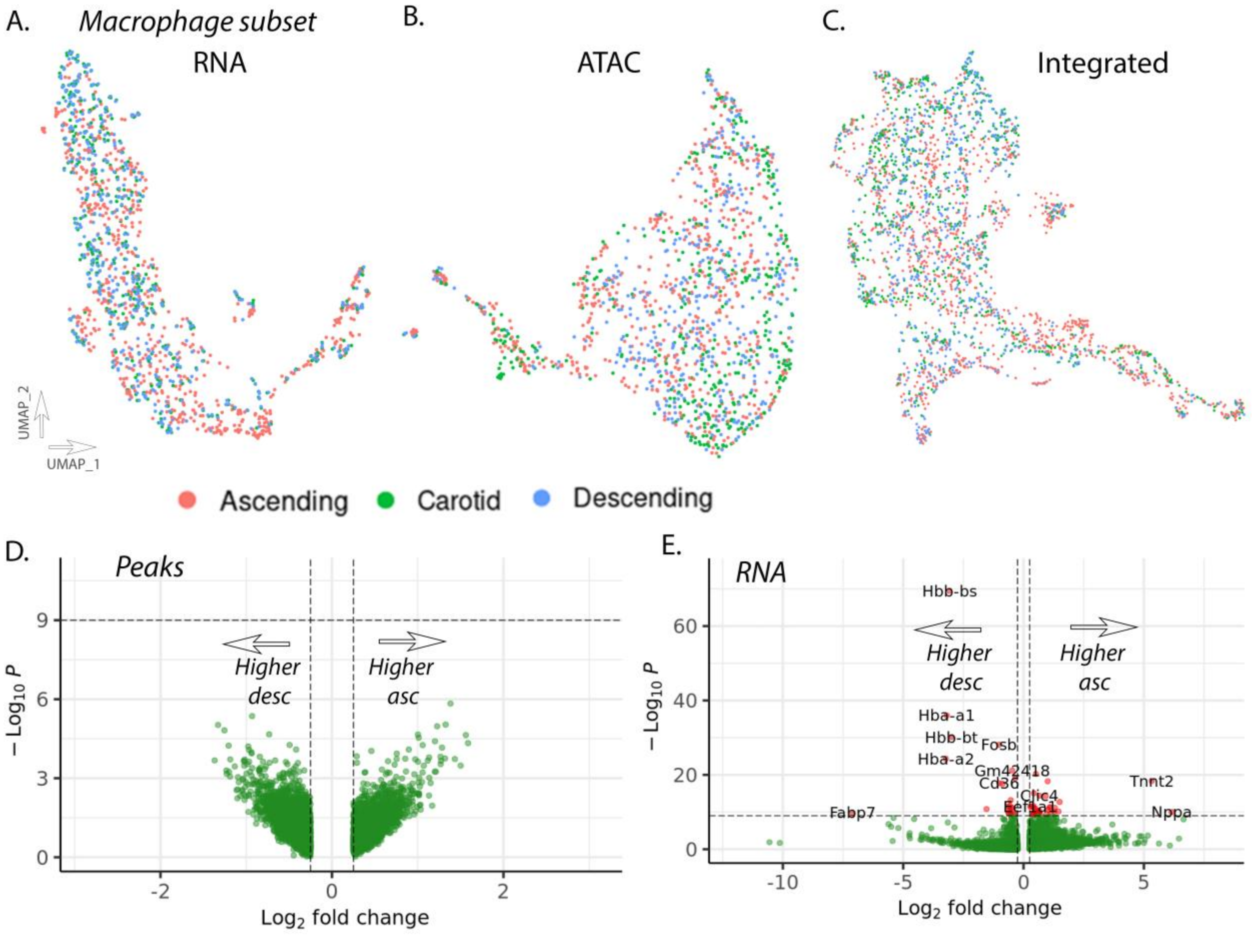


**Supplemental Figure 3. Single cell transcriptomic and epigenomic analysis of macrophage cells reveals notable homogeneity between vascular sites.** UMAP visualization of macrophage cell subset for RNA (A), ATAC (B), and integrated (C) datasets. Volcano plots for differential peak accessibility (D) and RNA expression (E).

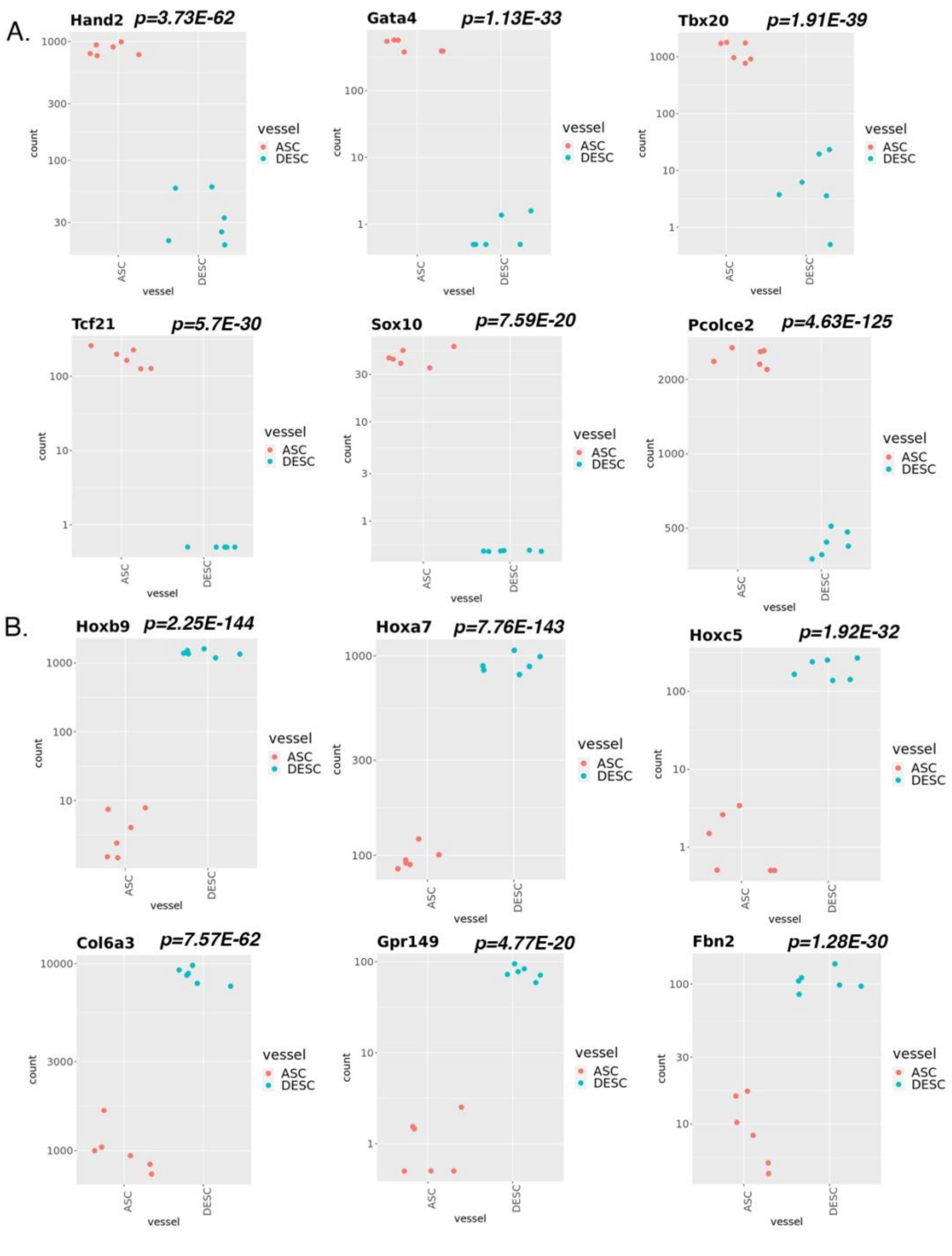


**Supplemental Figure 4.** RNA expression from primary adventitial fibroblast in vitro culture highlighting genes with increased expression in ascending fibroblasts including *Hand2*, *Gata4*, *Tbx20*, *Tcf21*, *Sox10*, and *Pcolce2* (A), and increased expression in descending fibroblasts including *Hoxb9*, *Hoxa7*, *Hoxc5*, *Col6a3*, *Gpr149*, and *Fbn2* (B). Y-axis represents normalized read counts by DESeq2 shown on log10 scale. P-value for comparison shown for each gene.

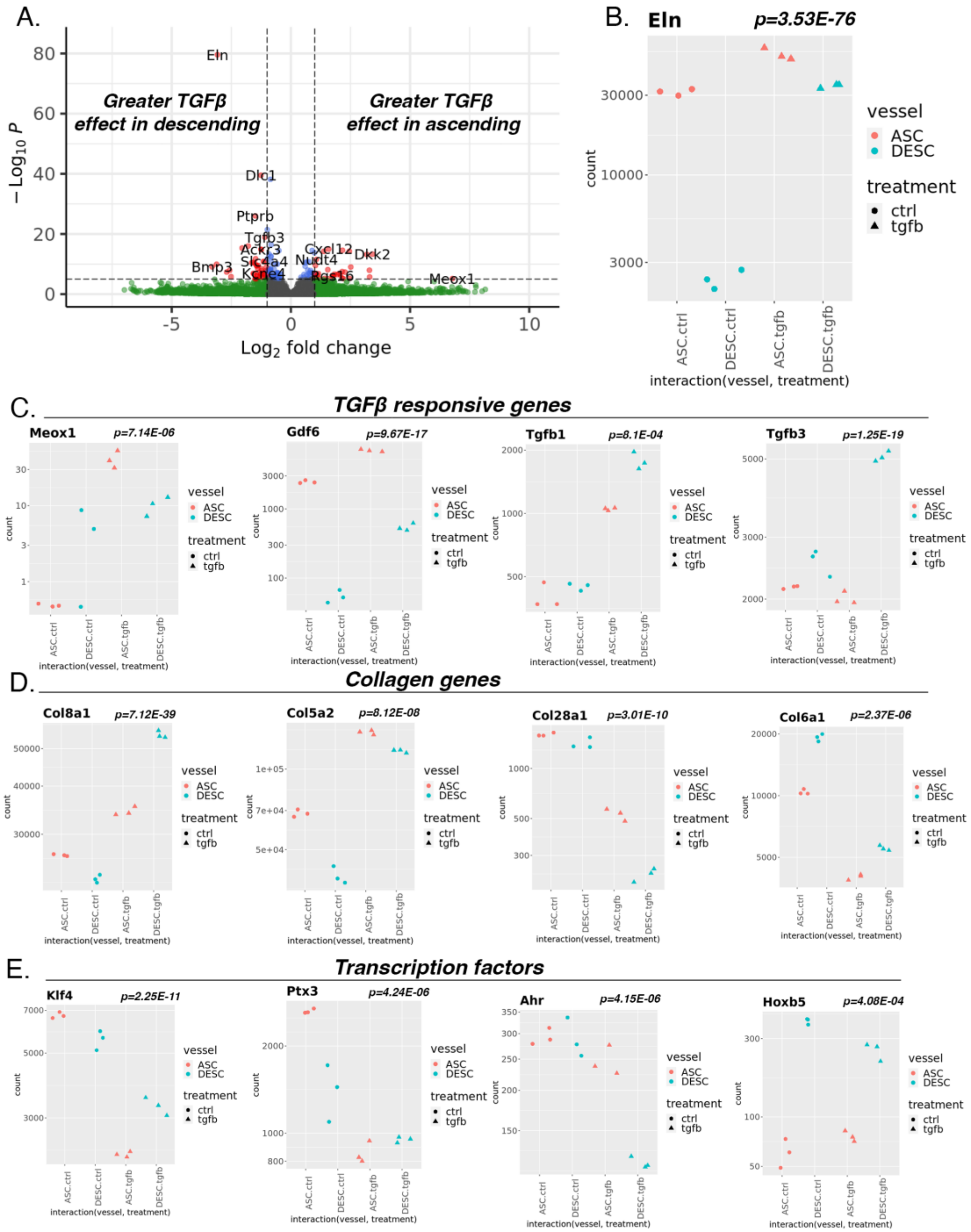


**Supplemental Figure 5.** Volcano plot of interaction analysis of RNAseq data revealing genes with differential response to TGFβ by vascular site (log2FC > 0 represents greater effect TGFβ in ascending; log2FC < 0 represents greater effect TGFβ in descending) (A). RNAseq expression for top interaction gene *Eln* (B). RNAseq expression for TGFβ responsive genes *Meox1*, *Gdf6*, *Tgfb1*, and *Tgfb3* (C) collagen genes *Col8a1*, *Col5a2*, *Col28a1*, *Col6a1* (D) and transcription factors *Klf4*, *Ptx3*, *Ahr*, and *Hoxb5* (E). Y-axis represents normalized read counts by DESeq2 shown on log10 scale. P-value for interaction shown for each gene.

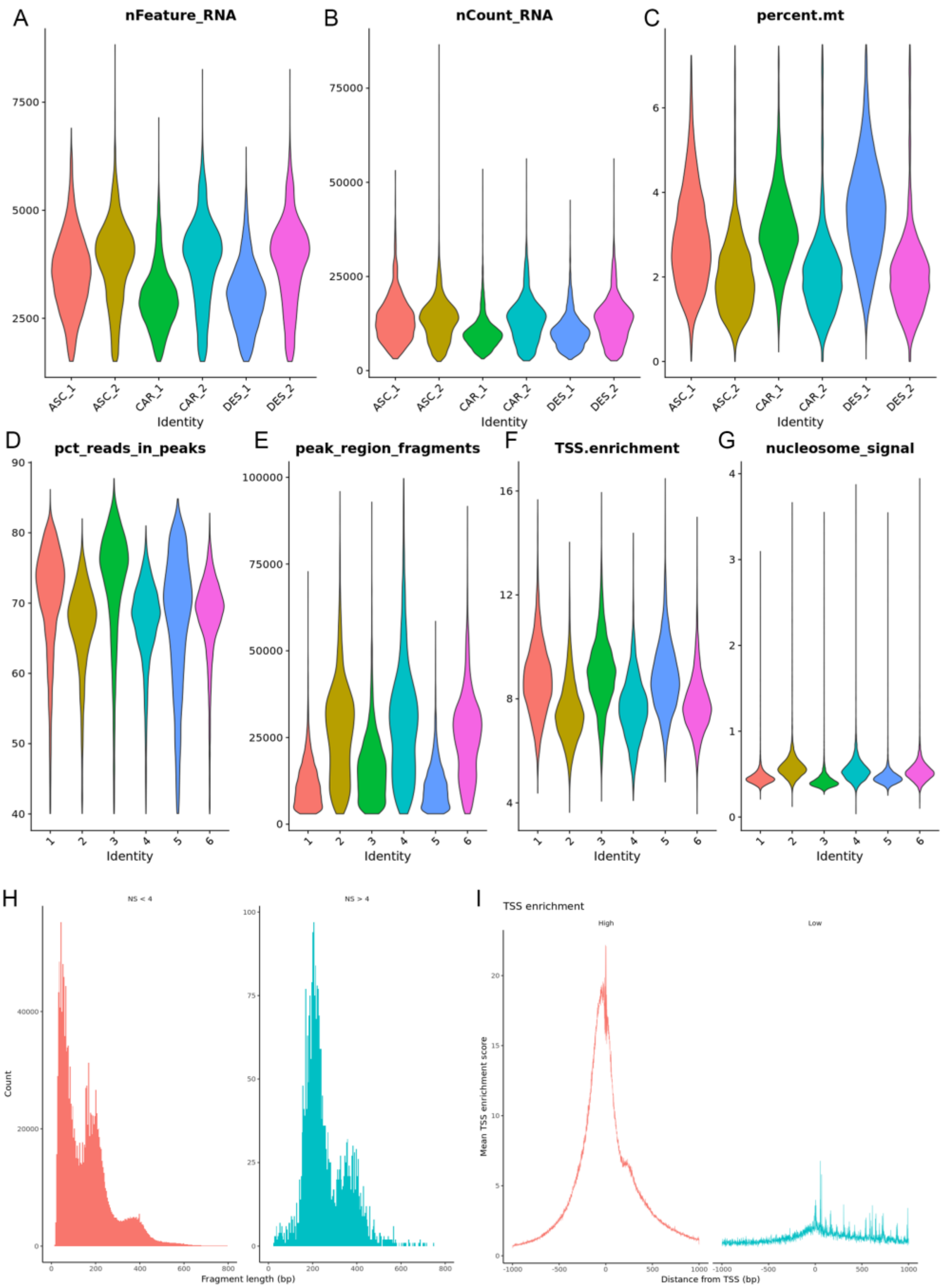


**Supplemental Figure 6.** QC metrics for scRNAseq and scATACseq datasets. For scRNAseq datasets, violin plots for nFeature_RNA (A), nCount_RNA (B), and percent.mt (C) across 6 captures. For scATACseq datasets, violin plots for pct_reads_in_peaks (D), peak_region_fragments (E), TSS.enrichment (F), and nucleosome_signal (G). Fragment histogram of region chr1-1-10000000 in combined aggr dataset based on low and high nucleosome signal (NS <4, left, NS >4, right) (H). TSS plot for combined aggr dataset based on high and low TSS enrichment (TSS >2, left, TSS <2, right).
